# TRPV4 calcium-permeable channel contributes to valve stiffening in aortic stenosis

**DOI:** 10.1101/2024.11.07.622108

**Authors:** Bidisha Dutta, Suneha G. Rahaman, Pritha Mukherjee, Shaik O. Rahaman

## Abstract

Aortic valve stenosis (AVS) is a progressive disease marked by fibrosis, inflammation, calcification, and stiffening of the aortic valve leaflets, leading to disrupted blood flow and left ventricular pressure overload. AVS can result in heart failure and death within 2 to 5 years if left untreated, highlighting its high mortality rate. Understanding the molecular mechanisms of AVS is essential for developing noninvasive treatments. Emerging data suggest that extracellular and intracellular matrix stiffness influences gene expression, inflammation, and cell differentiation. Myofibroblast activation of valvular interstitial cells (VICs) along with excess extracellular matrix (ECM) accumulation and remodeling are primary drivers of AVS progression. Inflammation also plays a critical role, with macrophages accumulating in valve leaflets from AVS patients, promoting inflammation, activating VICs, and synthesizing and remodeling the ECM.

Our lab and others have reported that macrophage and fibroblast activities, including migration, inflammatory gene expression, and myofibroblast activation, are sensitive to matrix stiffness, indicating that valve leaflet stiffening may regulate AVS progression via a cellular stiffness sensor. Our published work shows that mechanosensitive Ca^2+^-permeable transient receptor potential vanilloid 4 (TRPV4) channels regulate fibrosis in other organs and control macrophage and fibroblast activation, suggesting TRPV4 as the potential stiffness sensor in AVS. This implies that fibrosis and tissue stiffening may reinforce each other, creating a vicious cycle in AVS development, with VICs and macrophages playing central roles. Here, we identify the cellular stiffness sensor mediating the link between stiffness and AVS development using human aortic valve tissues, a murine model of aortic valve stenosis, and atomic force microscopy analysis.

## Introduction, Results, and Discussion

Aortic valve stenosis (AVS) is a progressive disease marked by fibrosis, inflammation, calcification, and stiffening of the aortic valve leaflets, leading to disrupted blood flow and left ventricular pressure overload (1, 2). Symptomatic AVS can result in heart failure and death within 2 to 5 years if left untreated, highlighting its high mortality rate (1, 2). Understanding the molecular mechanisms of AVS is essential for developing noninvasive treatments. Emerging data suggest that extracellular and intracellular matrix stiffness influences gene expression, inflammation, and cell differentiation (3, 4). Myofibroblast activation of valvular interstitial cells (VICs) along with excess extracellular matrix (ECM) accumulation and remodeling are primary drivers of AVS progression (1, 2). Inflammation also plays a critical role, with macrophages accumulating in valve leaflets from AVS patients, promoting inflammation, activating VICs, and synthesizing and remodeling the ECM (1, 2).

Our lab and others have reported that macrophage and fibroblast activities, including migration, inflammatory gene expression, and myofibroblast activation, are sensitive to matrix stiffness, indicating that valve leaflet stiffening may regulate AVS progression via a cellular stiffness sensor (3, 4). Our published work shows that mechanosensitive Ca^2+^-permeable transient receptor potential vanilloid 4 (TRPV4) channels regulate fibrosis in other organs and control macrophage and fibroblast activation, suggesting TRPV4 as the potential stiffness sensor in AVS (4, 5). This implies that fibrosis and tissue stiffening may reinforce each other, creating a vicious cycle in AVS development, with VICs and macrophages playing central roles. Here, we identify the cellular stiffness sensor mediating the link between stiffness and AVS development using human aortic valve tissues, a murine model of aortic valve stenosis, and atomic force microscopy analysis.

We presented clinical parameters of AVS patients and healthy subjects in Figure A. To detect TRPV4, CD68, and α-smooth muscle actin (α-SMA) proteins in human AVS valve tissues, we obtained 5 μm sections of aortic valve biopsies from AVS patients (n = 5) and healthy subjects (n = 2) (OriGene; Rockville, MD). Double immunofluorescence staining was performed on these sections to identify α-SMA-positive (α-SMA^+^) myofibroblasts and CD68-positive (CD68^+^) macrophages expressing TRPV4 proteins (TRPV4^+^). The immunofluorescence intensity was quantified as integrated density. TRPV4 activation is known to promote α-SMA expression, a marker of myofibroblast differentiation (4). Our lab’s previous work also shows TRPV4’s role in macrophage accumulation in fibrotic skin tissues (5). Based on this, we hypothesized that TRPV4 drives myofibroblast differentiation and macrophage accumulation in AVS. Our results showed an increased presence of α-SMA^+^ myofibroblasts and CD68^+^ macrophages in AVS valves compared to healthy controls, with both cell types coexpressing TRPV4 (Figure A). We observed a 10-fold increase in TRPV4, CD68, and α-SMA expression in AVS valves versus healthy controls (Figure A). Additionally, we found a strong correlation between TRPV4 and CD68 or α-SMA expression, suggesting TRPV4^+^-myofibroblasts and -macrophages are present in human AVS valve tissues (Figure A). To corroborate these findings, Western blot analysis confirmed an increased expression of TRPV4 protein in human stenotic samples compared to normal controls (Figure B).

**Figure.**
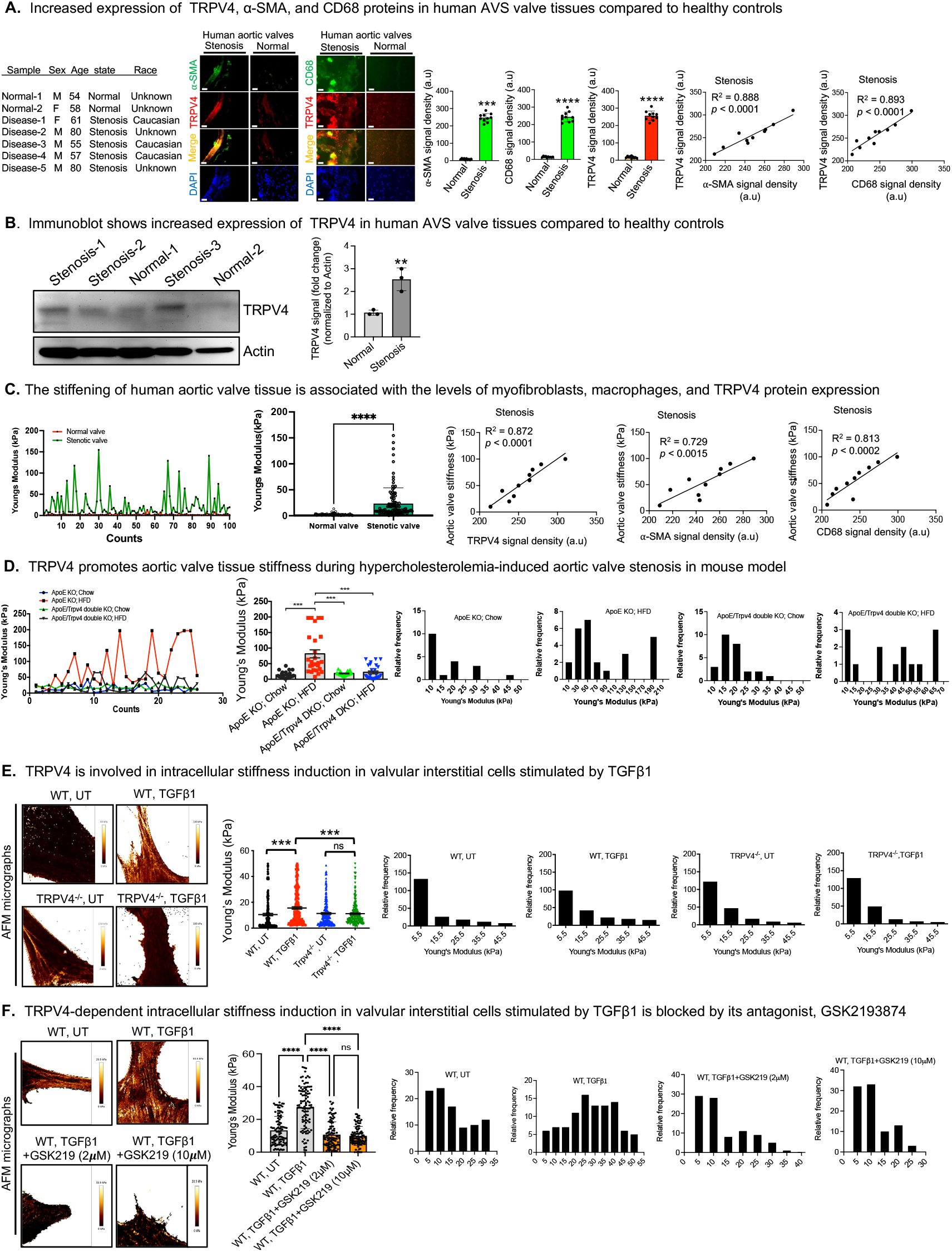
TRPV4 regulates matrix stiffening in AVS. **A**. Clinical parameters of subjects; Immunofluorescence staining of valve sections from AVS-patients or healthy-subjects to identify α-SMA^+^ myofibroblasts (anti-CD68; 1:100; Sigma-Aldrich) or CD68^+^ macrophages (anti-CD68; 1:100; Bio-Rad) expressing TRPV4 proteins (anti-TRPV4; 1:100; Alomone, Israel). Quantitation of results show expression of α-SMA, CD68, and TRPV4 (n = 5 sections/condition; \*\*\**p* < 0.001, \*\*\*\**p* < 0.0001, ANOVA); Scatterplot with linear regression analysis shows TRPV4 signals relative to α-SMA or CD68 abundance in five AVS-valve samples. **B**. Immunoblots show expression of TRPV4 and actin in stenotic and normal control tissues. Bar graph shows quantification of the data from three blots. \*\**p* < 0.01, t-test. **C**. Plot profile of Young’s modulus (YM) of the dataset (stenotic-valve vs. normal-valve) generated by AFM analysis. Quantification of YM of the dataset. n = 2-5 tissue samples/group and 50 force curves/sample. \*\*\*\**p* < 0.0001, ANOVA. Scatterplot with linear regression analysis shows the stenotic aortic valve stiffness relative to α-SMA, TRPV4, or CD68 abundance in five AVS-valve samples. **D**. Plot profile of YM of the dataset (ApoE^-/-^ vs. ApoE^-/-^Trpv4^-/-^ mice; HFD or chow for 6 months) generated by AFM analysis. n = 5 tissue samples/group and 30 force curves/sample; Quantification of YM of the dataset. n = 5 tissue samples/group and 30 force curves/sample. \*\*\**p* < 0.001, ANOVA; Representative frequency distributions of force curves. **E**. AFM micrographs to represent variations in stiffness in WT and TRPV4 KO VICs (untreated (UT) or TGFβ1 (5 ng/ml) for 48 h); Quantification of YM of the dataset generated by AFM analysis. n = 50 force curves/sample. \*\*\**p* < 0.001, ANOVA; Representative frequency distributions of force curves. **F**. AFM micrographs to represent variations in stiffness in WT VICs (untreated (UT) or TGFβ1 (5 ng/ml) with or without GSK2193874 (GSK219) for 27 h); Quantification of YM of the dataset generated by AFM analysis. n = 100 force curves/sample. \*\*\*\**p* < 0.0001, ns: not significant, ANOVA; Representative frequency distributions of force curves.

To test whether valve tissue stiffening is associated with human AVS, we measured aortic valve tissue stiffness (Young’s modulus) using atomic force microscopy (AFM) (5). Five-micrometer aortic valve sections from AVS patients and controls were analyzed to obtain force curves. We found valve tissue stiffness to be 12-fold higher in AVS valve tissues compared to healthy valves, indicating increased stiffness in AVS valves (Figure C). Additionally, analysis showed a strong correlation between tissue stiffness and levels of myofibroblasts (α-SMA^+^), macrophages (CD68^+^), and TRPV4 proteins, suggesting a role for these cells in tissue stiffening in AVS (Figure C).

To test whether TRPV4 regulates aortic valve stiffness in vivo in AVS mouse model, we measured aortic valve stiffness using AFM. Ten-micrometer aortic root sections from hypercholesterolemic ApoE^-/-^ and ApoE^-/-^Trpv4^-/-^ mice (C57BL/6, both genders, 6-8 weeks) were analyzed to obtain force curves. We found a 7-fold increase in aortic valve stiffness in ApoE^-/-^ mice compared to ApoE^-/-^Trpv4^-/-^ mice after 6 months on a high-fat diet (HFD) (21% fat, 0.15% cholesterol; Harlan Teklad) or control chow, suggesting TRPV4’s role in regulating aortic valve stiffness during AVS progression (Figure D). AVS development involves valve stiffening due to extracellular matrix remodeling and cellular stiffness changes, though the exact mechanism remains unknown.

To test whether TRPV4 regulates cellular stiffness in VICs, the primary cellular component of aortic valve leaflets, we measured stiffness in mouse primary VICs using AFM. We found significantly increased stiffness in wildtype-VICs compared to TRPV4 KO VICs stimulated by TGFβ1, a profibrotic factor linked to organ fibrosis and stiffening, suggesting TRPV4’s role in regulating VIC stiffness during AVS progression (Figure E). To support these findings, AFM analysis demonstrated a reduction in stiffness in VICs treated with the TRPV4 antagonist GSK2193874 compared to vehicle controls (Figure F).

TRPV4 regulates cytoskeletal dynamics through interactions with small GTPases like RhoA and Rac1, affecting actin remodeling, cell migration, differentiation, and cell shape. These processes could significantly impact AVS development and progression. We suggest that RhoA-mediated TRPV4 modulation may play a key regulatory role in VICs. Although TRPV4-driven RhoA signaling activation has been observed in fibroblasts and trabecular meshwork cells, its role in VICs remains unexplored in the context of AVS. Activation of RhoA signaling by TRPV4 may influence VIC differentiation into myofibroblasts and promote matrix remodeling, thereby contributing to mechanical stress responses and fibrotic processes. This, in turn, can create a positive feedback loop that reinforces matrix stiffening and TRPV4 activation, potentially driving AVS development and progression.

Altogether, our data indicates that TRPV4 contributes to matrix stiffening in AVS. VICs may not be the only source of myofibroblasts in AVS, as other cell types, including endothelial cells and pericytes, may differentiate into myofibroblasts during fibrosis. Electrophysiological experiments on human tissues to demonstrate TRPV4 activity and its modulation by matrix stiffness could provide more direct evidence of TRPV4 channel activation and its potential role in the development and progression of AVS. This approach would offer valuable insights into how mechanical cues are translated into AVS-related pathological changes through TRPV4 signaling. Further studies are needed to determine how TRPV4 regulates fibrosis and matrix stiffening in AVS and to explore the potential of targeting TRPV4 therapeutically to ameliorate AVS.

## Materials and Methods

### Mice and stenosis model

Dr. Makato Suzuki (Jichi Medical University, Tochigi, Japan) initially generated TRPV4 knockout (KO) mice on a C57BL/6 background, which were subsequently obtained from Dr. David X. Zhang (Medical College of Wisconsin, Milwaukee, WI). ApoE KO mice were acquired from the Jackson Laboratory, and ApoE:TRPV4 double knockout (DKO) mice on a C57BL/6 background were developed by the Rahaman lab. All mice were housed and bred in a temperature- and humidity-controlled, germ-free environment with ad libitum access to food and water. Experimental procedures adhered to the guidelines of the Institutional Animal Care and Use Committee and were approved by the University of Maryland review committee. For the stenosis model, ApoE KO and ApoE:TRPV4 DKO mice were fed a high-fat diet (HFD; 21% fat, 0.15% cholesterol; Harlan Teklad) or a control chow diet for six months.

### Valvular interstitial cell (VIC) culture and maintenance

We harvested VICs from mouse hearts using a protocol adapted from JOVE Journal (6). VICs were isolated from WT and TRPV4 KO mice by dissecting heart valve areas. Valve leaflets were treated with collagenase II and incubated at 37°C for 1 hour, followed by vigorous vortexing to remove endothelial cells. The leaflets were minced and incubated in collagenase II for an additional 15 minutes, then pipetted to release cells. Cells were washed with PBS, resuspended in DMEM with 10% FBS, and plated on collagen-coated plates. Fresh medium was provided every 48 hours. For passaging, 0.25% trypsin-EDTA was used, and cells were frozen in 10% DMSO with 90% FBS. Experiments used cells at passage 3-8, with TGFβ1 treatment performed in DMEM containing 1% BSA.

### Immunofluorescence staining

To detect TRPV4, CD68, and α-smooth muscle actin (α-SMA) proteins in human AVS tissues, we obtained 5 μm sections from aortic valve biopsies of AVS patients (n = 5) and healthy controls (n = 2) (OriGene; Rockville, MD). Double immunofluorescence staining was performed to identify α-SMA-positive (α-SMA+) myofibroblasts and CD68-positive (CD68+) macrophages expressing TRPV4 (TRPV4+). Immunofluorescence intensity was measured as integrated density. Tissue sections were stained overnight at 4°C with anti-CD68, anti-TRPV4, and anti-SMA antibodies (1:100) to detect TRPV4+ macrophages and myofibroblasts. Alexa Fluor 488- and 594-conjugated anti-rabbit or anti-mouse IgG (1:200) were used for secondary staining, and DAPI was applied to stain cell nuclei.

### Western blot analysis

For Western blot analysis of TRPV4 and actin levels, tissue lysates were prepared from aortic valve biopsies of AVS patients (n = 3) and healthy controls (n = 2). Pierce RIPA buffer (cat# 89900) and Halt protease and phosphatase inhibitor (cat# 78442) from Thermo Fisher Scientific were used for lysis. Protein extracts were separated on 10% SDS-PAGE gels and probed with anti-TRPV4 (1:1000) and anti-actin (1:2000) antibodies. HRP-conjugated secondary IgGs were used for visualization, and blots were analyzed with the UVP BioSpectrum system.

### Atomic force microscopy

We utilized the JPK Nano Wizard 4 atomic force microscope (AFM) (Bruker Nano GmbH, Berlin, Germany) to assess stiffness in aortic valve tissues (human and mouse) and valvular interstitial cells (VICs) (5). For tissue samples, 7-10 μm-thick aortic valve sections were mounted on glass slides and submerged in PBS to record individual force spectroscopy curves (F-D curves) in contact mode for each sample. Measurements were performed using CP-qp-CONT-BSG-B-5 colloidal probes (sQube) with a 30 kHz resonance frequency in air, a spring constant of 0.1 N/m, gold-coated on the detector side, and a 10 μm borosilicate glass sphere. Parameters included a setpoint of 1 nN, extension speed of 10 μm/s, and Z length of 10 μm to ensure appropriate F-D curves, recording 50 F-D curves per sample across various regions. The Hertz model for spherical probes was applied to calculate Young’s modulus (E). Data visualization, including histograms and plots, was conducted using GraphPad Prism.

For VIC stiffness measurements on living condition, AFM was operated in advanced force-spectroscopy based mode, referred to as Quantitative Imaging (QI™), which enables simultaneous nanomechanical characterization and imaging. VICs were seeded on poly-D-lysine-coated glass-bottom dishes (FD35PDL, WPI) and treated with or without 5 ng/mL TGFβ1 and GSK2193874 (2 and 10 μM, Sigma Aldrich) for 24 hours. For measurements, cells were maintained in PBS at 37°C with 5% CO_2_. QI maps of 128 × 128 pixels were obtained. For these measurements, qpBioAC-CI-CB2 cantilevers (Nanosensors) with a 50 kHz resonance frequency in air, a spring constant of 0.1 N/m, partial gold coating on the detector side, and a 30 nm-radius quartz probe were used. We imaged a total of 10 cells per condition, scanning 3-4 areas per cell with an imaging setpoint of 0.5 nN.

### Sources of funding

This work was supported by an NIH (R01EB024556) grant to Shaik O. Rahaman.

## Data availability

All data generated or analyzed during this study are included in this article.

## References

1. Goody PR, Hosen MR, Christmann D, Niepmann ST, Zietzer A, Adam M, Bönner F, Zimmer S, Nickenig G, Jansen F. Aortic valve stenosis: From basic mechanisms to novel therapeutic targets. Arterioscler Thromb Vasc Biol. 2020;(April):885–900.

2. Bruschi G, Maloberti A, Sormani P, Colombo G, Nava S, Vallerio P., Casadei F, Bruno J, Moreo A, Merlanti B, Russo CD, Oliva F, Klugmann S, Giannattasio C. Arterial Stiffness in Aortic Stenosis: Relationship with Severity and Echocardiographic Procedures Response. High Blood Press Cardiovasc Prev. 2017;24(1):19–27.

3. Humphrey JD, Dufresne ER, and Schwartz MA (2014) Mechanotransduction and extracellular matrix homeostasis. Nat. Rev. Mol. Cell Biol 15, 802–812

4. Rahaman SO, Grove LM, Paruchuri S, Southern BD, Abraham S, Niese KA, Scheraga RG, Ghosh S, Thodeti CK, Zhang DX, Moran MM, Schilling WP, Tschumperlin DJ, Olman MA. TRPV4 mediates myofibroblast differentiation and pulmonary fibrosis in mice. J Clin Invest. 2014;124(12):5225–5238.

5. Goswami R, Arya RK, Sharma S, Dutta B, Stamov DR, Zhu X, Rahaman SO. Mechanosensing by TRPV4 mediates stiffness-induced foreign body response and giant cell formation. Sci Signaling. 2021, Nov 2: 14 (707): eabd4077.

6. Bouchareb, R., Lebeche, D. Isolation of Mouse Interstitial Valve Cells to Study the Calcification of the Aortic Valve In Vitro. J. Vis. Exp. (171), e62419, doi:10.3791/62419 (2021).

